# The molecular basis of Kale domestication: Transcription profiling of leaves and meristems provides new insights into the evolution of a *Brassica oleracea* vegetative morphotype

**DOI:** 10.1101/2020.11.25.398347

**Authors:** Tatiana Arias, Chad Niederhuth, Paula McSteen, J. Chris Pires

## Abstract

Morphotypes of *Brassica oleracea* are the result of a dynamic interaction between genes that regulate the transition between vegetative and reproductive stages and those that regulate leaf morphology and plant architecture. In kales ornate leaves, delayed flowering, and nutritional quality are some of the characters potentially selected by humans during domestication.

We used a combination of developmental studies and transcriptomics to understand the vegetative domestication syndrome of kale. To identify candidate genes that are responsible for the evolution of domestic kale we searched for transcriptome-wide differences among three vegetative *B. oleracea* morphotypes. RNAseq experiments were used to understand the global pattern of expressed genes during one single phase of development in kale, cabbage and the rapid cycling kale line TO1000.

We identified gene expression patterns that differ among morphotypes, and estimate the contribution of morphotype-specific gene expression that sets kale apart (3958 differentially expressed genes). Differentially expressed genes that regulate the vegetative to reproductive transition were abundant in all morphotypes. Genes involved in leaf morphology, plan architecture, defense and nutrition were differentially expressed in kale.

RNA-Seq experiments allow the discovery of novel candidate genes involved in the kale domestication syndrome. We identified candidate genes differentially expressed in kale that could be responsible for variation in flowering times, taste and herbivore defense, variation in leaf morphology, plant architecture, and nutritional value. Understanding candidate genes responsible for kale domestication is of importance to ultimately improve Cole crop production.

## INTRODUCTION

Domestication is the process of selection used by humans to adapt wild plants to cultivation. During this process, a set of recurrent characters is acquired across a wide diversity of crops (known as the “domestication syndrome”) (Hammer, 1984; Doebley, 1992). Recurrent traits observed in domesticated plants include: loss of seed shattering, changes in seed size, loss of photoperiod sensitivity, and changes in plant physiology and architecture (Doebley, 1992; Harlam, 1992; Allaby 2014). Here we explore the molecular basis behind the poorly defined vegetative domestication syndrome of kale, a *Brassica oleracea* morphotype, the leaves of which are consumed by humans and are also used as fodder. The domestication syndrome of kale includes: apical dominance, ornate leaf patterns, the capacity to delay flower formation and maintain a vegetative state producing a higher yield of the edible portion, nutritional value of leaves, and the ability to defend against herbivores that also give the leaves a unique pungent taste.

The Cole crops (*Brassica oleracea*) are perennial plants native to Europe and the Mediterranean and include a range of domesticated and wild varieties (Snogerup, 1980; Tsunoda, 1980; Purugganan et al., 2000; Allender et al., 2007; Maggioni et al. 2010). Each crop type is distinguished by domestication traits not commonly found among the wild populations (Maggioni et al., 2010). They include dwarf plants with a main stem that has highly compressed internodes (cabbages); plants with an elongated main stem in which the lateral branches are highly compressed due to very short internodes (Brussels sprouts); plants with proliferation of floral meristems (broccoli) or proliferation of aborted floral meristems (cauliflower); and plants with swollen stems (kohlrabi, Marrow-stem kale) or ornate leaf patterns (kales) (Hodgkin, 1995; Lan and Paterson, 2000). It is not so easy to define which domestication traits are common to all of the above-mentioned types, as opposed to the wild species. At early stages in domestication, plants must have been selected for less bitter-tasting, less fibrous, thicker stems, and more succulent storage organs (Maggioni et al., 2010).

Research has provided evidence that polyploidy and genetic redundancy contribute significantly to the phenotypic variation observed among these crops (Lukens et al., 2004; Paterson, 2005). The genetic basis underlying the phenotypic diversity present in Cole crops has been addressed only for some of these morphotypes. The cauliflower phenotype of *Arabidopsis* is comparable to the inflorescence development in the cauliflower *B. oleracea* morphotype. It has been shown that the gene *CAULIFLOWER* is responsible for the phenotype observed in both species (Kempin et al., 1995; Carr & Irish, 1997; Purugganan et al., 2000; Smith and King, 2000). Also, a large QTL for the curd of cauliflower has been found (Lan and Paterson, 2000) indicating that 86 genes are controlling eight curd-related traits, demonstrating this phenotype is under complex genetic control. Other studies have found and confirmed *BoAP-a1* to be involved in curd formation in cauliflower (Anthony et al., 1995; Carr and Irish, 1997; Gao et al., 2007) and have shown that morphologies in this cultivar are under complex genetic control (Sebastian et al., 2002). The molecular bases of broccoli have also been proposed as involving changes in three codons in the *BoCAL* and *BoAP* genes (Carr and Irish, 1997; Lowman and Purugganan, 1999). A total of five QTLs have been detected for leaf traits in Brussel sprouts, but no significant QTLs have been found for axillary buds (Sebastian et al., 2002). Head formation in cabbage has been suggested as additively inherited (Tanaka et al., 2009). In kale, a recent study suggest that the lobed-leaf trait is quantitatively inherited (Ren et al., 2019).

Apical dominance is manifest in kale varieties, involving selection for erect plants with fewer side branches, and more compact plants, which allows more plants to fit into each unit of cultivated soil (Deham et al., 2020). In kale, leaves are green to purple and do not form a compact head like in cabbage. Crop varieties have surprising variation in leaf size, leaf type, and margin curliness. Some also have an over-proliferation of leaf blade tissues. Kale leaf curliness, blade texture, and tissue overgrowth are some of the more variable morphological characters among varieties.

Leafy kales are considered the earliest cultivated brassicas, originally used for both livestock feed and human consumption (Maggioni et al., 2010) due to leaves and floral buds with great nutritional value (Velasco et al., 2011). Kale leaves also have antioxidative properties, and have been shown to reduce cholesterol levels (Velasco et al., 2007; Soengas et al., 2012). Their high content of beta-carotene, vitamin K, vitamin C, and calcium make it one of the most nutritious vegetables available for human consumption (Stewarta and McDougalla, 2012). Kale is a source of two carotenoids: lutein and zeaxanthin, and contains sulforaphane, a compound suggested to have anti-cancer properties (Velasco et al., 2007). Kale leaves also have mustard oils produced from glucosinolates. These natural chemicals most likely contribute to plant defense against pests and diseases, and impart a pungent flavor property characteristic of all cruciferous vegetables (Sonderby et al., 2010; Carmona et al., 2011). Definitions of domestication syndrome in kale include traits affecting cultivation, harvesting, and cooking – and its uses – including food, medicines, toxins and fibres (Deham et al., 2020).

Here we used a combination of developmental studies and transcriptomes to understand the vegetative domestication syndrome of kale. Comparisons of kale morphotypes with the “rapid-cycling” morphotype TO1000 and cabbage allow us to identified transcriptome-wide differences that set kale apart. RNA-seq of meristems and immature leaves for the *B. oleracea* vegetative morphotypes, cabbage, kale, and TO1000 reveal gene expression patterns unique to kale and associated developmental and metabolic processes. This allowed us to identify a set of candidate genes we suggest may be important in the kale domestication syndrome.

## MATERIALS AND METHODS

### Plant Material and Experimental Design

Single accessions of cabbage (*B. oleracea* var. *sabauda*), kale (*B. oleracea* var. *viridis*) and the rapid cycling TO1000 (*B. oleracea* var. *alboglabra*) were grown in an environmental chamber under uniform conditions: 10hr day (light 7AM to 5PM, Dark 5:01PM to 6:59AM daily) and watered every other day. Morphotypes were planted in a completely randomized block design. Initially, two seeds per pot were planted, then only one was picked (32 plants per morphotype total). A random number generator was used to move plants and flats around every other day. Structural features of the meristems and developing leaves were observed using an Environmental Scanning Electron Microscopy (ESEM) and light microscopy. Images were processed with Adobe (San Jose, California, USA) Photoshop, version 7.0.

Tissues were collected 24 days after seeds were planted. The upper portion of the stems, including immature leaves, and apical and lateral meristems were flash frozen in liquid nitrogen. The assumption was made that genes involved in developing different morphologies should be expressed in younger leaves, shoots, and meristems. Three biological replicates per morphotype were collected, each replicate consisting of pooled tissue from eight plants.

### RNA Extraction and cDNA Synthesis

Lysis and homogenization of fresh soft tissue were performed before RNA extraction; PureLink Micro-to-Midi Total RNA Purification System (Invitrogen, Cat. No 12183-018, Carlsbad, California, USA) was used. RNA extraction was done using the Purelink RNA Mini Kit (Ambion, Cat. No12183-018A, Carlsbad, California, USA). DNase was not used before proceeding with cDNA synthesis as recommended in the protocol. For cDNA synthesis MINT (Evrogen, Cat. No SK001, Moscow, Russia) was used, this kit enzymes synthesize full-length-enriched double stranded cDNA from a total polyA^+^ RNA. The cDNA was then amplified by 16 --21 cycles of polymerase chain reaction and purified using a PCR Purification Kit (Invitrogen K3100-01 Carlsbad, California, USA).

### Library Construction and Illumina Sequencing

Protocols previously published by Steele et al. (2012) were implemented here. End repair was performed on 400μl with 3μg of normalized ds-cDNA prior to ligating barcoding adapters for multiplexing, using NEB Prep kit E600L (New England Biolabs, Ipswich, Massachusetts, USA). Shearing was done using a Bioruptor® sonication device (Diagenode, Denville New Jersey, USA), at 4°C, continuously, for 15 min. total (7 mix) on high. Three replicates per morphotype were tagged with different adaptors and sequenced on a single lane of an Illumina GAIIx machine.

Oligonucleotides used as adapters were: Cabbage (1: ACGT, 2: GCTT, 3: TGCT), kale (1: TACT, 2:ATGT, 3:GTAT), TO1000 (1:CTGT, 2:AGCT, 3:TCAT). To prepare adapters, equal volumes were combined of each adapter at 100 μmol/L floating them in a beaker of boiling water for 30 min and snap cooling them on ice. Ligation products were run on a 2% low-melt agarose gel and size-selected for ~300 bp using an x-tracta Disposable Gel Extraction Tool (Scientific, Ocala Florida, USA). Fragments were enriched using PCR in 50 μL volumes containing 3 μL of ligation product, 20 μL of ddH2O, 25 μL master mix (from NEB kit), and 1 μL each of a 25 μmol/L solution of each forward and reverse primer (forward 5′-AAT GAT ACG GCG ACC ACC GAG ATC TAC ACT CTT TCC CTA CAC GAC GCT CTT CCG ATC* T-3′; reverse 5′-CAA GCA GAA GAC GGC ATA CGA GAT CGG TCT CGG CAT TCC TGC TGA ACC GCT CTT CCG ATC*-3′; both HPLC purified). Thermal cycle routine was as follows: 98°C for 30 s, followed by 15 cycles of 98°C for 10 s, 65°C for 30 s, and 72°C for 30 s, with a final extension step of 72°C for 5 min. Products were run on a 2% low-melt agarose gel and the target product was excised and purified with the tools described above. DNA was purified at each step during the end repair, adapter ligation, size selection and fragment enrichment with either a Gel Extraction kit (Qiagen, Germantown Maryland, USA) or a DNA QIAquick PCR Purification kit (Qiagen, Germantown Maryland, USA). The three replicates per morphotype were ran on one third of a lane with single-end 98-bp reads on an Illumina GAIIx Genome Analyzer using the Illumina Cluster Generation Kit v2-GA II, Cycle Sequencing Kit v3 (Illumina, San Diego, USA), and image analysis using Illumina RTA 1.4.15.0 (Illumina, San Diego, USA) at University of Missouri DNA Core.

### Data quality control and preprocessing, alignment and differential expression analysis

Reads were first trimmed for the end quality using Cutadapt v2.4 (Martin et al., 2011) and read quality assessed using FastQC (Andrews, 2010). These were then aligned against the *Brassica oleracea* TO1000 genome (Parkin et al., 2014) downloaded from EnsemblPlants 44 (Kersey et al., 2018) with two passes of STAR v2.7.1a (Dobin et al., 2013). In the first pass, reads for each sample were aligned for splice-junction discovery, in addition to those junctions provided in the TO1000 annotations. Splice junctions from each sample were then provided as a list for a second final alignment and read counts calculated for each gene using STAR. Mapping statistics are provided in Supplementary Table 1. Raw sequencing data and unnormalized read counts are publicly available through the Gene Expression Omnibus accession GSE149483. Read counts were imported into R v3.6.3 (Team R) and differential expression was determined using DESeq2 v1.26.0 (Love et al., 2014). Specifically, read counts were normalized, with the TO1000 samples as reference, and fitted to a parametric model. We looked for differentially expressed genes (DEGs) for each pairwise comparison (kale vs TO1000, kale vs cabbage, and cabbage vs TO1000). Comparisons were done to ultimately separate unique kale-morphotype DEGs. Since each pairwise comparison increases the potential number of false positives, these were combined into a single table and the false discovery rate (FDR) (Benjamini and Hochberg, 1995) was adjusted for the entire table. Genes with an FDR < 0.05 and Log-fold change > 1 or < −1 were considered as differentially expressed genes (DEGs). Plots were made using the R packages ggplot2 (Wickham, 2009), pheatmap (Kolde, 2019), and a VennDiagram (Delhomme, 2018). Transcription-associated proteins (TAPs) were identified using the approach described for the construction of PlnTFDB v3.0 (http://plntfdb.bio.uni-potsdam.de) (Riaño-Pachón et al., 2007; Pérez-Rodríguez et al., 2010). All code for bioinformatic analysis and original figures pdfs and tables are available on GitHub (https://github.com/niederhuth/The-molecular-basis-of-Kale-domestication).

### Gene annotation, GO, and KEGG analysis

Gene ontology (GO) terms (Ashburner et al., 2000) are not currently available for the TO1000 assembly. In order to annotate TO1000 gene models with GO terms, TO1000 protein sequences were Blasted (blastp) against protein sequences from UniProtKB Swiss-Prot (release 2019_05) (Consortium 2018) using Diamond (Buchfink et al., 2014), with an e-value cutoff of 1e^−5^ and a max of 1 hit per query. UniProt IDs were then used to extract GO terms and mapped to the corresponding TO1000 gene model. In total, 35,422 genes (out of 59,225 total, ~59.8%) had a GO term assigned. GO term enrichment was performed using topGO v 2.38.1 (Alexa and Rahnenfuhrer, 2019) and GO.db v3.10.0 (Carlson, 2019) with the parent-child algorithm (Grossman et al., 2007) and Fisher’s Exact Test. A minimum node size of 5 was required, i.e. each GO term required a minimum of 5 genes mapping to it in ordered to be considered. P-values were adjusted for FDR and considered significant with an adjusted p-value < 0.05.

## RESULTS

### Kale leaf development

Developmental transitions and morphological changes were observed in two kale varieties (curly and non-curly kale) and the rapid cycling TO1000 for twenty-two days (Fig. 1–2). A week after planting all morphotypes had germinated; at first only the cotyledons were observed. Curly-kale grew faster than non-curly kale and TO1000. Using Environmental Scanning Electron Microscopy (ESEM), tiny true leaves with lobed margins and abundant trichomes were observed for curly-kale at five days (Fig. 2N), while non-curly kale and TO1000 had not developed lobes or leaves at this point (Fig. 2M, O). Eight-day-old curly-kale first immature leaves were seen emerging from between the cotyledons and covered by trichomes (Fig. 2B). Ten-days after germination curly-kale seedlings had abundant trichomes and highly lobed leaf margins; leaf shape was elliptic to obovate and leaves had red venation and red petioles (Fig. 2E). In comparison to the kale varieties, TO1000 seedlings had taller hypocotyls and erect stems. Leaf shape was oblanceolate and leaf margins were sinuate (Fig. 2D). Twelve-day-old curly-kale produce ectopic meristems in the leaf margins and more sporadically in the leaf blade of mature leaves; microscopic scales surrounded these ectopic meristems (Fig. 2Q). Ectopic meristems located at the tip of each lobe had a vascular bundle inserted to them and were always subtended by a trichome. Nineteen-day-old curly-kale and TO1000 had four leaves, the fourth one small; while non-curly kale has three leaves and the third one small. Twenty to twenty-two day-old curly-kale leaves develop more lobes and less trichomes with time. TO1000 leaves were thinner with sinuate margins. In summary, we observed kale margins were more lobed than TO1000 (Fig. 2).

**Figure 1.**
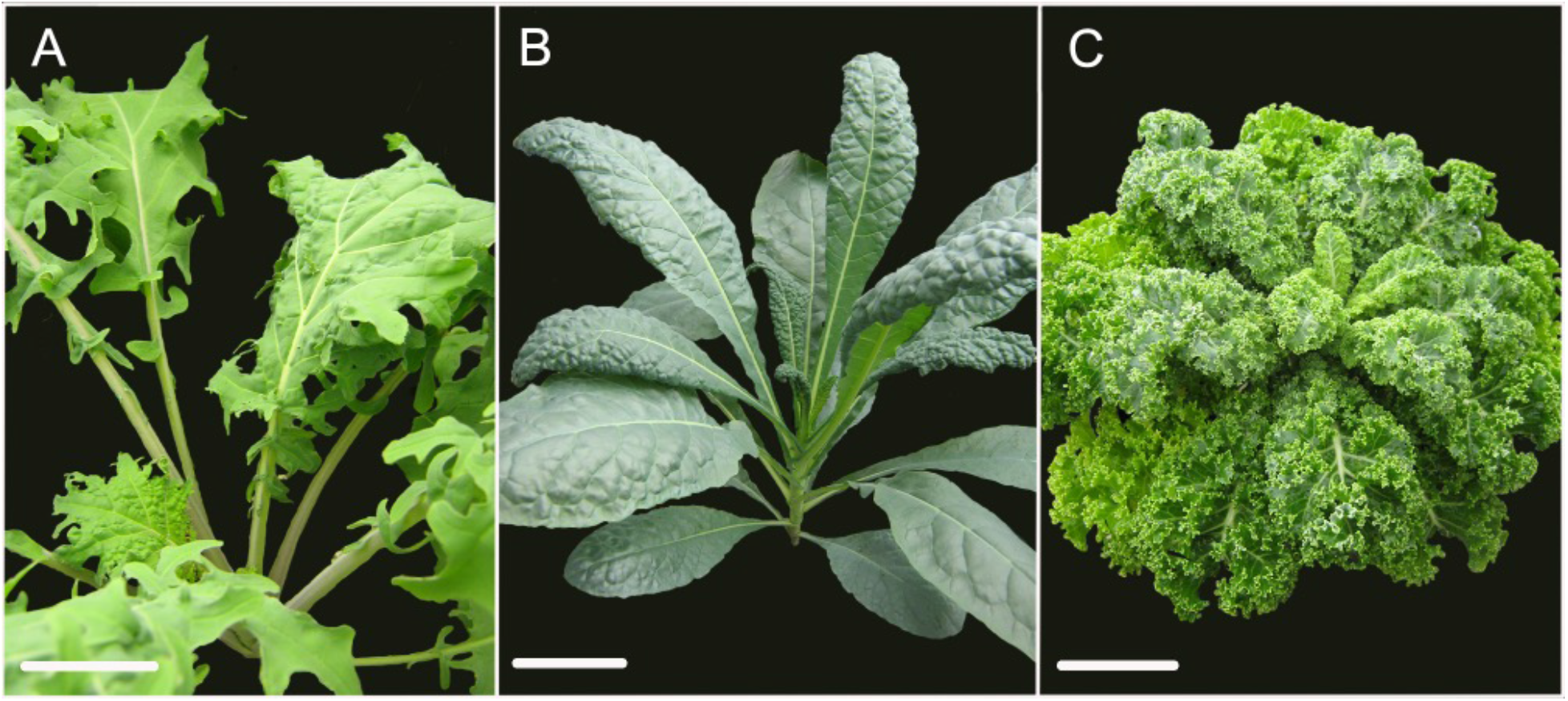
Comparison of mature leaf morphologies within kale varieties. (A) red winter kale (curly kale) with lobed leaf margins, (B) Lacinato Nero Toscana kale (non-curly kale) with crenulate leaf margins, (C) parsley kale with crisped leaf margins. Scale bar = 10 cm

**Figure 2.**
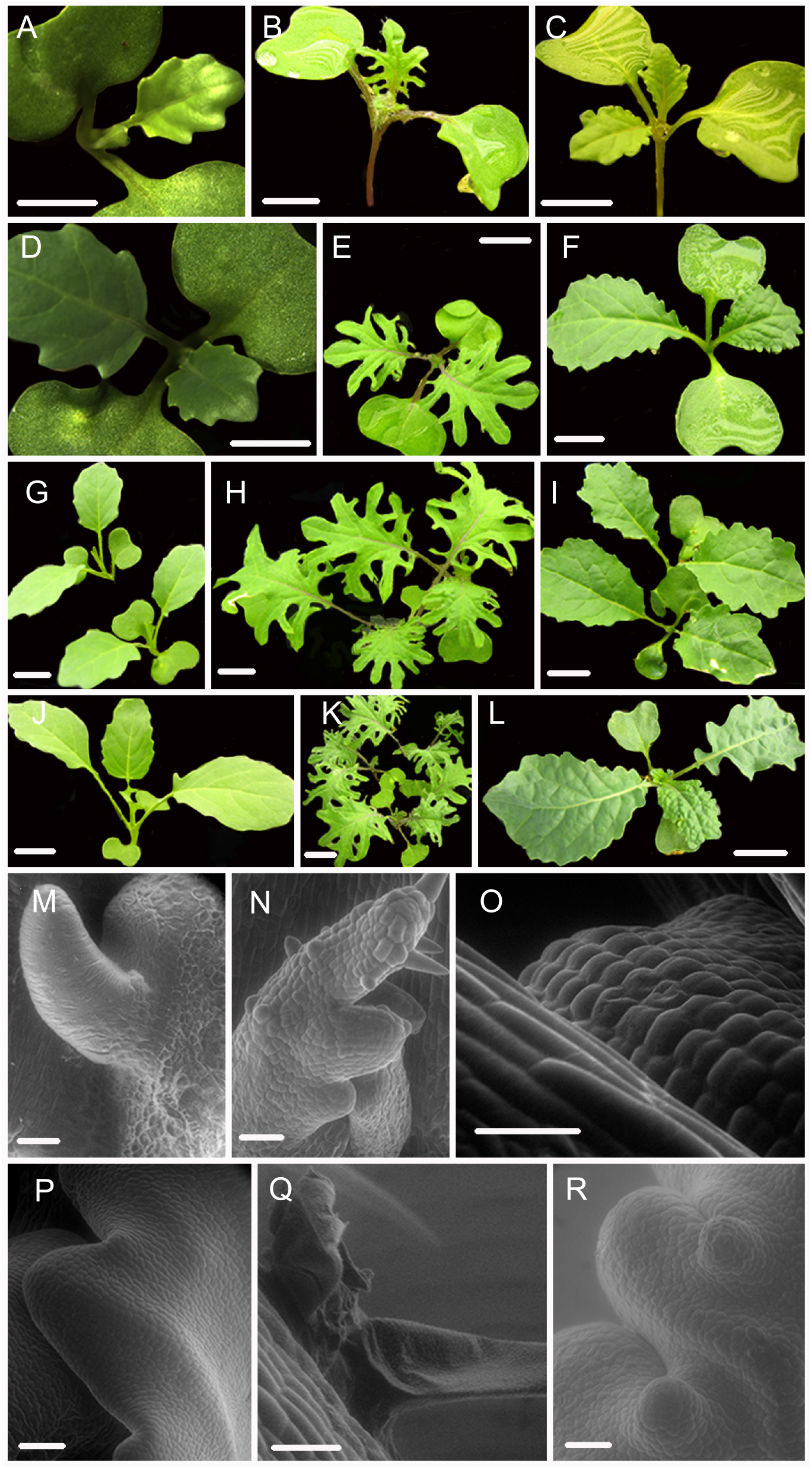
Leaf shape development in different Kale varieties. A-L: progressive development of the different Kale varieties and the rapid cycling TO1000 control (stereoscope and regular pics), scale bar = 1 cm. M-R: Scanning Electron Microscopy (SEM) of apical meristem, primordial leaves and details of lobes in different kale varieties and TO1000 control, scale bar = 5 microns. A, D, G, J: leaf development series for TO1000. B, E, H, K: leaf development series for curly kale (red winter kale). C, F, I, L: leaf development series for non-curly kale. M, P: SEM of TO1000 leaf primordia and margin formation detail. N: SEM of curly kale leaf primordial. O: SEM of non-curly kale (Lacinato Nero Toscana kale) leaf primordia. Q: SEM of ectopic growing of leaf blade in mature leaves subtended by a trichome. R: Non-curly kale (Lacinato Nero Toscana kale) margin formation detail.

### RNA-seq and differential gene expression

To identify genes that might be associated with domestication traits including leaf development in curly-kale, RNA-seq was used to assess transcriptional differences between curly-kale, cabbage, and TO1000. Three biological replicates each were sequenced, however, one of the cabbage replicates was not used for assembly because very few reads of low quality were obtained after sequencing. Between 9 to 25 million trimmed reads were obtained per sample, with ~78-89% reads mapping uniquely to the TO1000 *B. oleracea* genome (Suppl. Table 1). After normalization with DESeq2, hierarchical clustering and principle component analysis showed that samples were grouped by morphotype, with greater variance between morphotypes than between replicates (Suppl. Fig. 1).

Differentially expressed genes (DEGs) were called for all pair-wise comparisons between morphotypes (kale vs TO1000, kale vs cabbage, cabbage vs TO1000), with the aim of identifying genes that were differentially expressed in kale compared to other morphotypes. Genes were considered differentially expressed if they had a two-fold change in expression and a p-value < 0.05 after correction for multiple testing. In total, 8854 DEGs were identified between kale and TO1000 (4599 higher expressed in kale, 4255 lower expressed in kale), 8037 DEGs between kale and cabbage (4704 higher expressed in kale, 3333 lower expressed in kale), and 8189 DEGs between cabbage and TO1000 (3406 higher expressed in cabbage, 4783 lower expressed in cabbage) (Fig. 3A). There were a total of 3958 DEGs shared between the kale-TO1000 and kale-cabbage comparisons, 2960 found only in the kale comparisons and 998 found in all three comparisons (Fig. 3B).

**Figure 3.**
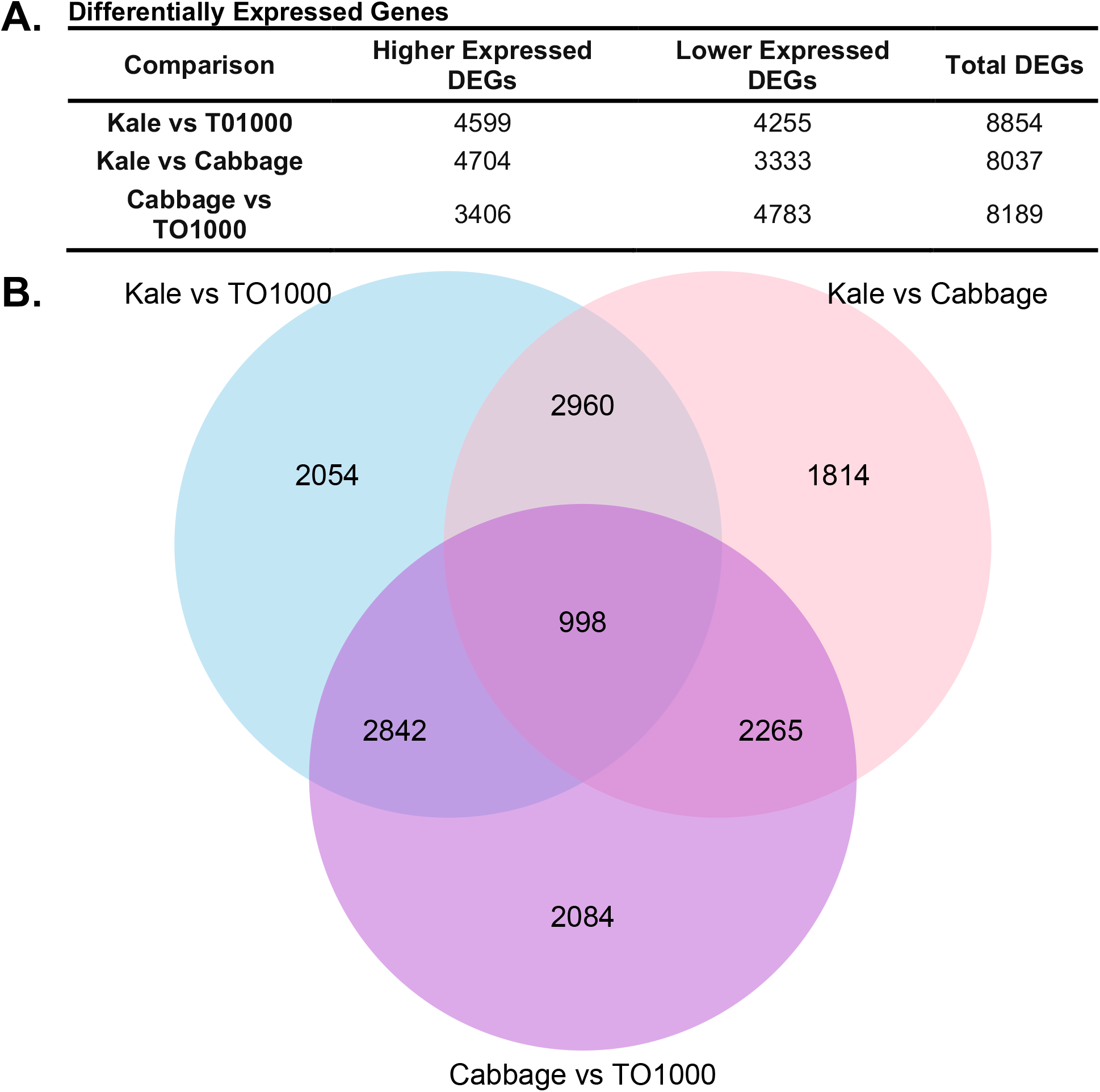
Number of genes differentially expressed in this study after comparing three different *Brassica oleracea* morphotypes (cabbage, kale and TO1000). A. Number of genes differentially expressed. B. Venn Diagram showing shared genes after doing all possible comparisons between three different *B. oleracea* morphotypes

### Polyploidy and its contribution to the kale phenotype

We explored how each of the three subgenomes resulting from an ancient whole genome triplication contributed to differential gene expression between morphotypes. Syntenic genes were significantly overrepresented amongst DEGs while non-syntenic genes were underrepresented in the kale-cabbage and cabbage-TO1000 comparisons, but not the kale-TO1000 comparisons (Fig. 4A). The gene content of the three sub-genomes of *B. oleracea* have been previously reconstructed (Parking et al., 2014) and using this, the representation of each sub-genome amongst DEGs was tested (Fig. 4C-D). The LF (least-fractionated) subgenome was underrepresented in kale-TO1000 comparison (odds ratio = 0.9162346. FDR-corrected p-value = 4.116097e-03) and overrepresented in the cabbage-TO1000 comparison (odds ratio = 1.0816227. FDR-corrected p-value = 9.948104e-03). The MF1 (more-fractionated 1) subgenome was significantly overrepresented for DEGs in all three comparisons (kale-TO1000: odds ratio = 1.0912796, FDR-corrected p-value < 1.103789e-02; kale-cabbage: odds ratio = 1.2091591, FDR-corrected p-value < 5.909977e-08; cabbage-TO1000: odds ratio = 1.1776233, FDR-corrected p-value = 2.913654e-06). The MF2 (more-fractionated 2) subgenome was overrepresented in the cabbage-TO1000 compariosn (odds ratio = 1.1118552. FDR-corrected p-value = 4.931485e-03).

**Figure 4.**
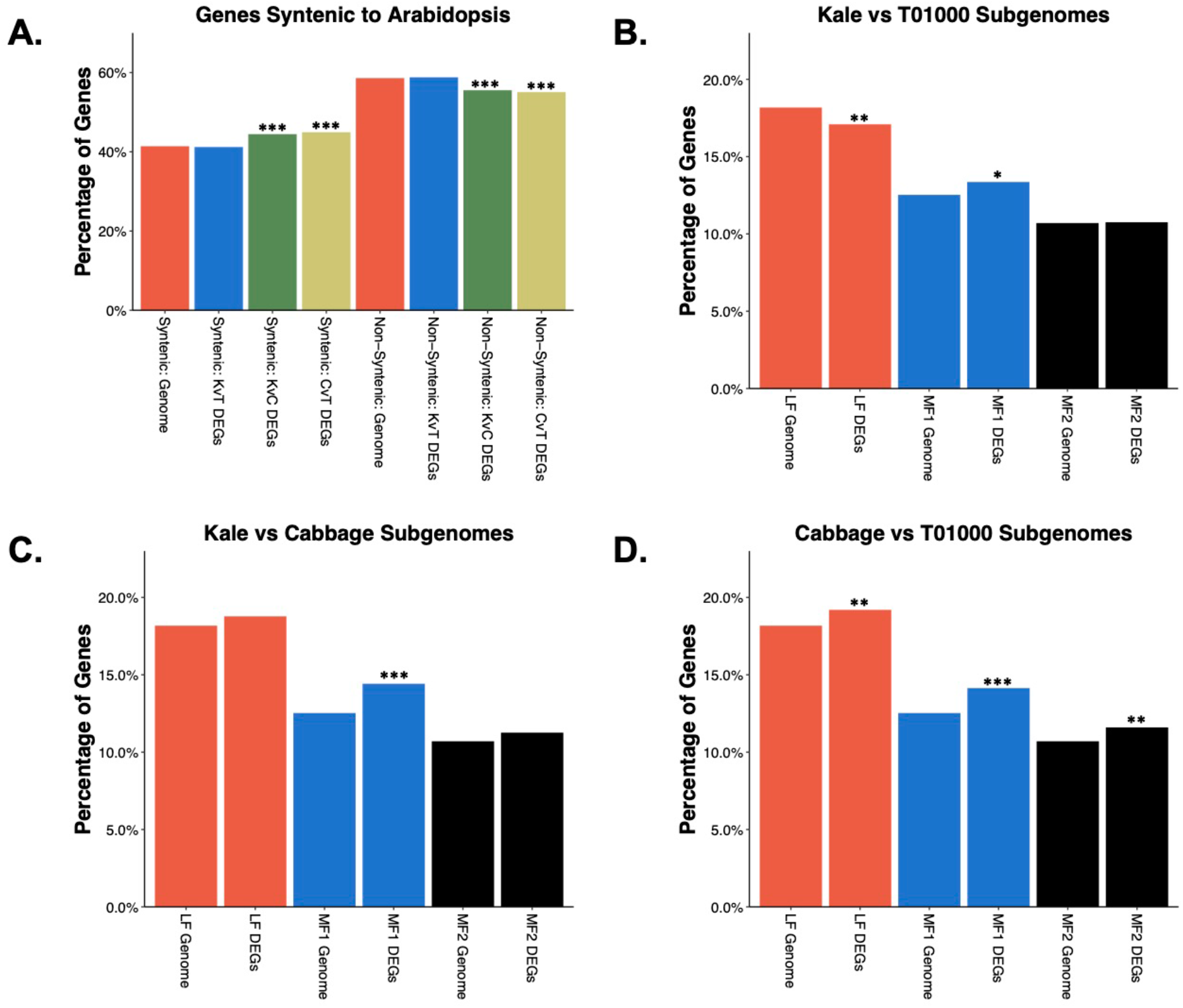
Syntenic genes from the three subgenomes are enriched in Differentially Expressed Genes (DEGs).

### Gene ontology and KEGG pathway enrichment analysis

DEGs for each pair-wise comparison and the list of DEGs shared between the two kale comparisons (3958 genes) were separated into lists of higher and lower-expressed genes and then tested for enrichment of GO (Gene Ontology) terms (Suppl. Tables 5-12) and KEGG (Kyoto Encyclopedia of Genes and Genomes) pathways (Suppl. Tables 13-20). We focus here specifically on those kale comparisons to better understand the kale domestication syndrome, in particular GO terms in the biological process (BP) category, as this includes developmental terms. GO terms and KEGG pathways related to the cabbage vs TO1000 comparison are provided as supplement (Suppl. Fig. 2).

#### Kale vs. TO1000

There were 91 (64 BP) and 17 (1 BP) GO terms enriched in the higher (Suppl. Table 5) and lower (Suppl. Table 6) expressed DEGs, respectively. Higher expressed genes were characterized by a number of metabolic processes: ‘lipid metabolism’, ‘nitrile metabolism,’ ‘sulfur compound metabolism’, benzene-containing compound metabolism’, ‘organic hydroxy compound biosynthesis’, and ‘carbohydrate derivative catabolism’. There were also multiple GO terms related to photosynthesis and plastid organization and environmental responses to both biotic and abiotic stresses. The only developmental terms enriched were related to senescence (Suppl. Fig. 2A). Lower expressed genes showed little enrichment, except for genes involved in ‘cellular nitrogen compound metabolism’ (Suppl. Figure 2B). Six KEGG pathways were enriched in higher expressed genes (Suppl. Table 13) and largely correspond to metabolic pathways identified in the GO term analysis. These include processes related to ‘photosynthesi-antenna proteins’, ‘fatty acid elongation’, ‘flavonoid biosynthesis’, ‘steroid biosynthesis’, ‘glyoxylate and dicarboxylate metabolism’, ‘other glycan degradation’, and ‘cyanoamino acid metabolism’. Lower expressed genes were enriched in ‘spliceosome’ and ‘circadian rhythm’ pathways (Suppl. Table 14).

#### Kale vs. cabbage

Higher expressed genes had 21 (15 BP) enriched GO terms (Suppl. Table 7), while lower expressed genes (Suppl. Table 8) had 26 (11 BP). Similar to the kale vs TO1000 comparison, there was enrichment for processes involved in ‘fatty acid derivative metabolism’, ‘nitrile metabolism’, and ‘photosynthesis’ (Suppl. Figure 2C). There was also enrichment for environmental responses, specifically responses to herbivores. Additionally, there was enrichment for processes in ‘carbon fixation’, ‘benzoate metabolism, ‘endosome organization’, and the ‘negative regulation of ion transport. Lower expressed genes (Suppl. Fig. 2D) include ‘glycosinolate metabolism’, ‘response to chitin’, and ‘gene expression’. ‘Sulfur compound metabolism’ was enriched in both comparisons, but was enriched amongst lower-expressed genes in the kale-cabbage comparison, while being enriched in higher-expressed genes in the kale-TO1000 comparison. These are again reflected in KEGG pathways where higher expressed genes were enriched for ‘other glycan degradation’ KEGG pathways (Suppl. Table 15). Lower expressed genes were enriched for ‘glucosinolate biosynthesis’ and ‘glutathione metabolism’ KEGG pathways (Suppl. Table 16).

#### Shared genes

Higher expressed genes were enriched for 32 (22 BP) GO terms (Suppl. Table 11 and Suppl. Fig. 2G). These again include multiple metabolic terms, especially related to ‘fatty acid derivative metabolism’, ‘nitrile metabolism’, ‘carbohydrate derivative catabolism’, and ‘benzoate metabolism’. Also similar to the individual comparisons, ‘photosynthesis’ and terms related to abiotic and biotic stresses, especially herbivores, were enriched. Unique to this subgroup was an enrichment for protein phosphorylation terms. Only four GO terms were enriched in the lower expressed genes and these were all in the cellular component GO term class (Suppl. Table 12). Higher expressed DEGs were enriched for ‘other glycan degradation’ and ‘Sesquiterpenoid and triterpenoid biosynthesis’ KEGG pathways. The former being also enriched in both of the individual comparisons. No KEGG pathways were enriched in the shared lower expressed DEGs (Supp. Tables 19 and 20).

### Candidate genes potentially involved in the domestication of kale

A set of candidate genes were identified based on their expression pattern (Fig. 5) and functions related to the suite of kale domestication syndrome characteristics. We further identified transcription factors (TF) families and found a total of 141 TF in a total of 3010 differentially expressed genes (Fig. 3B). Some of the most abundant included bHLH, ERF and MYB TFs (Suppl. Table 21).

**Figure 5.**
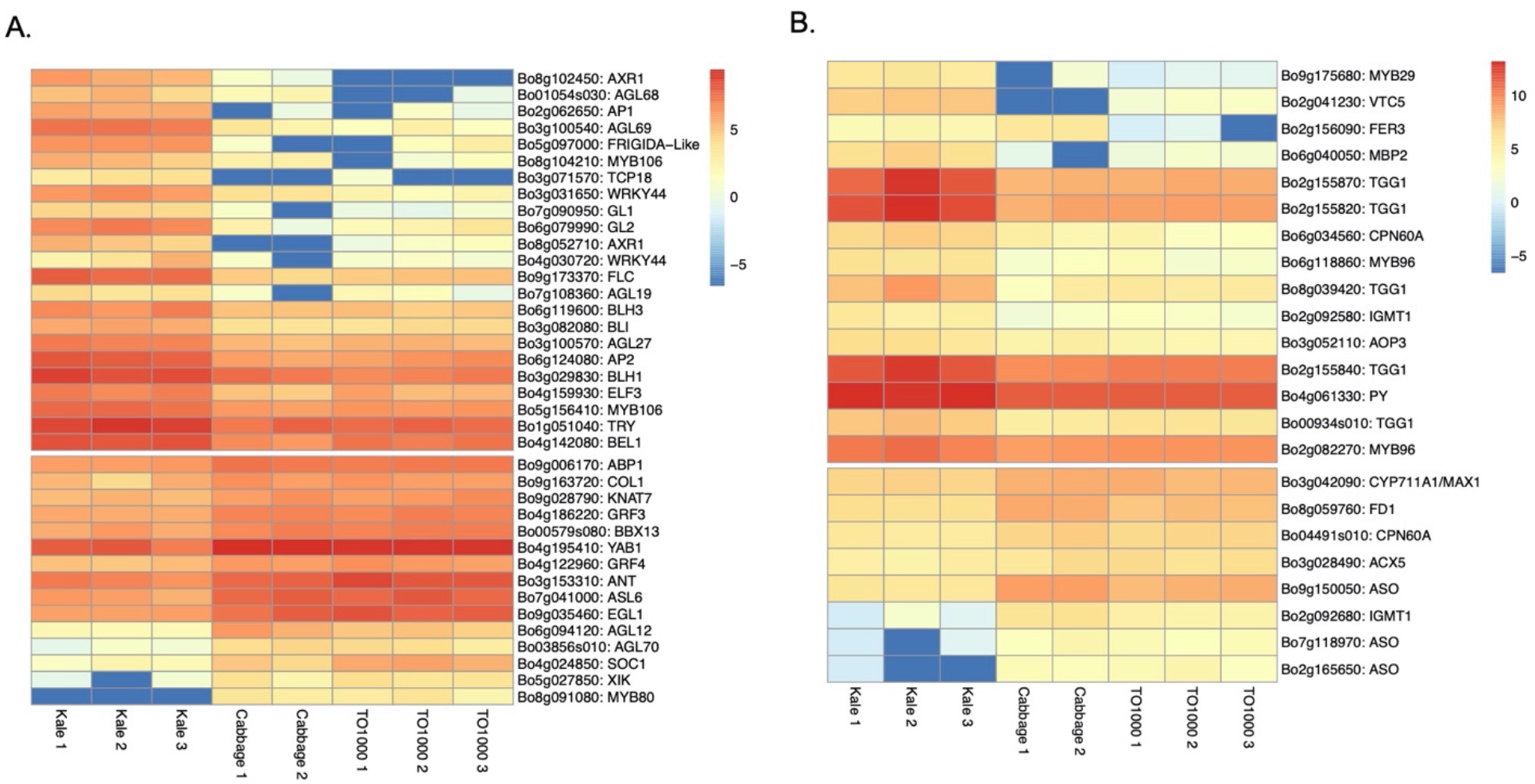
Heatmap of differentially expressed genes for curly kale (red winter kale). Up and down-regulated genes involved in A. plant and leaf development, and flowering B. plant defence responses and taste, nutrition.

#### Apical dominance and plant architecture

Several genes known to be involved in shoot grow inhibition were differentially expressed in kale tissues and upregulated, these include: an auxin-resistance *AXR1* (Bo8g102450, Bo8g052710) (Stirnberg et al., 1999), *TCP18* (Teosinte Branched 1-Like 1) (Bo3g071570), two copies of *REVOLUTA* (Bo9g130730, Bo7g021280) (Poza-Carrión et al., 2007), and the Beta-carotene isomerase D27 chloroplastic (Protein *DWARF-27* homolog) (Bo9g035190) (Waters et al., 2012). The More Axillary Branches gene *MAX1* (Bo3g042090) (Bennett et al., 2006) was also differentially expressed and downregulated in kale (Table 1; Fig. 5A).

**Table 1.**
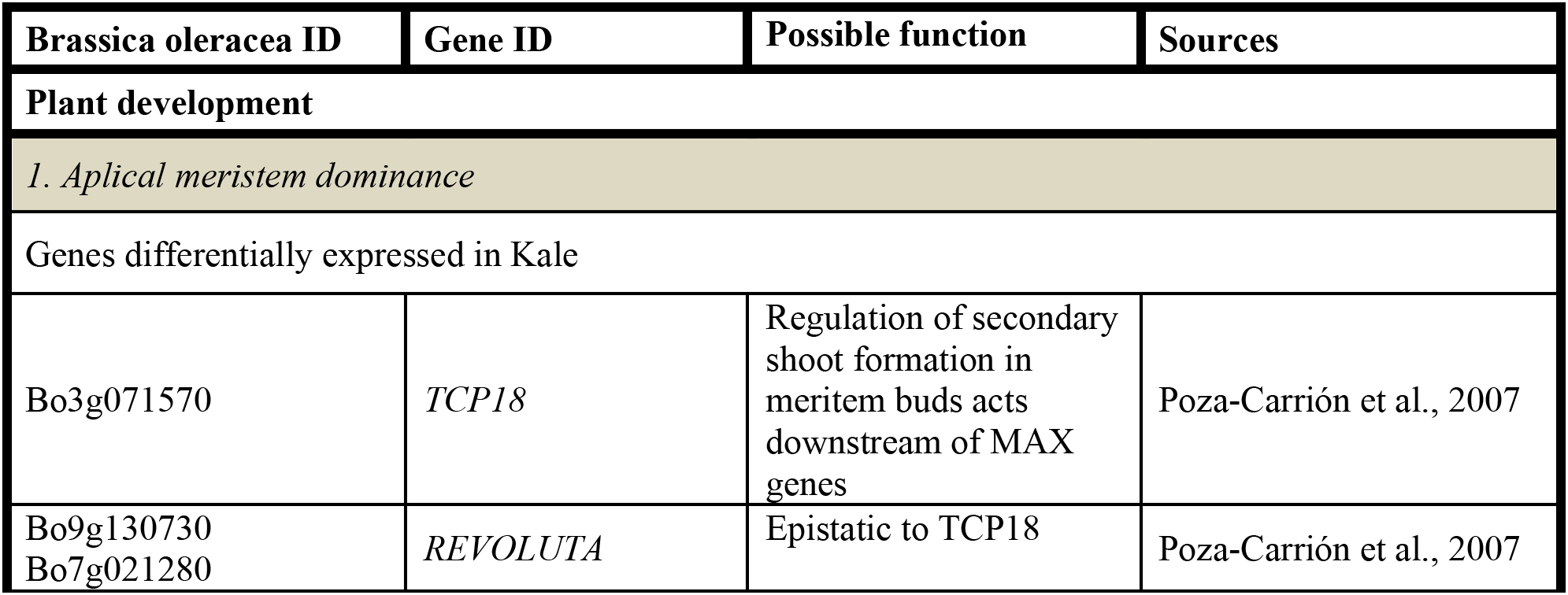
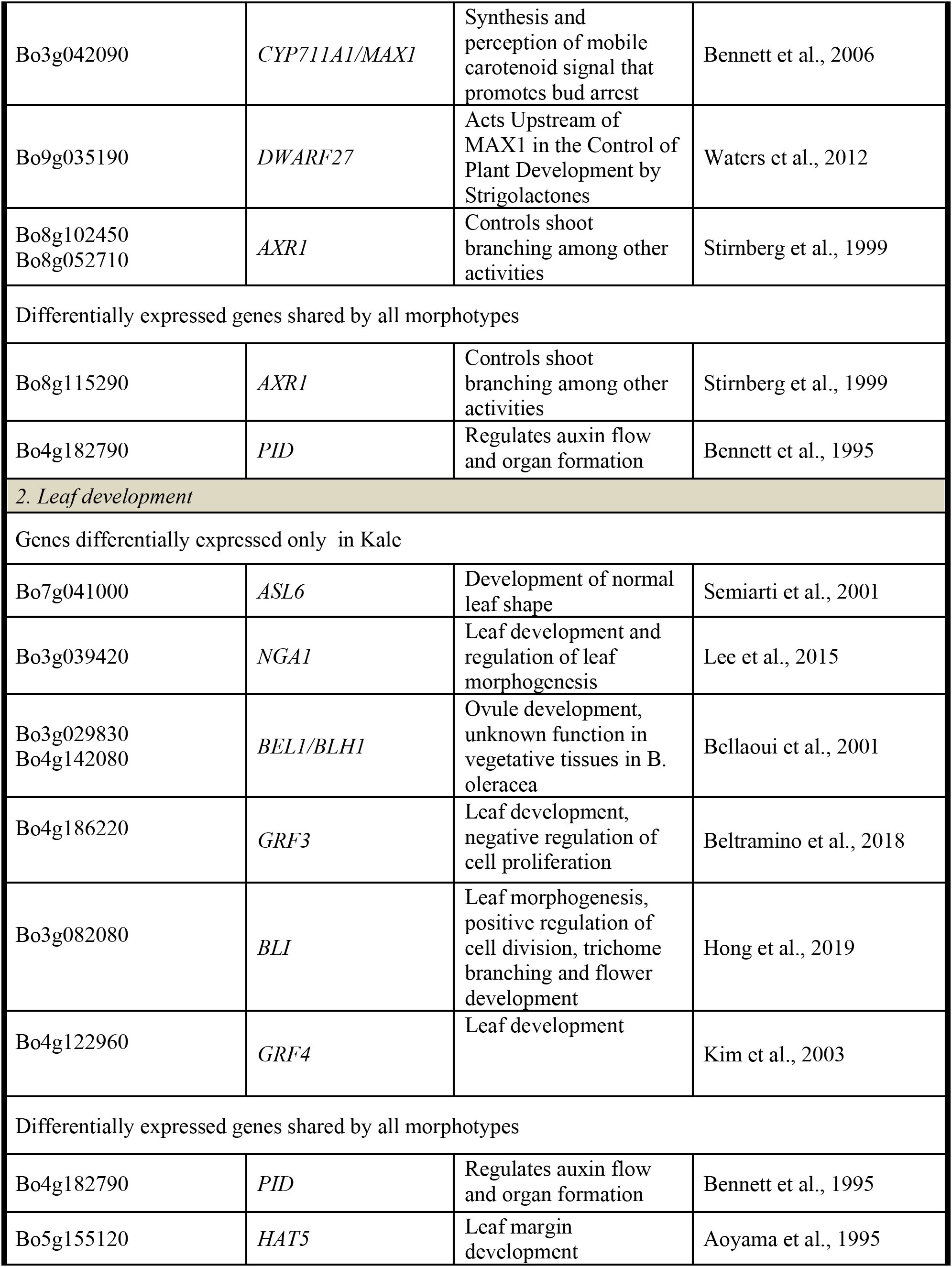
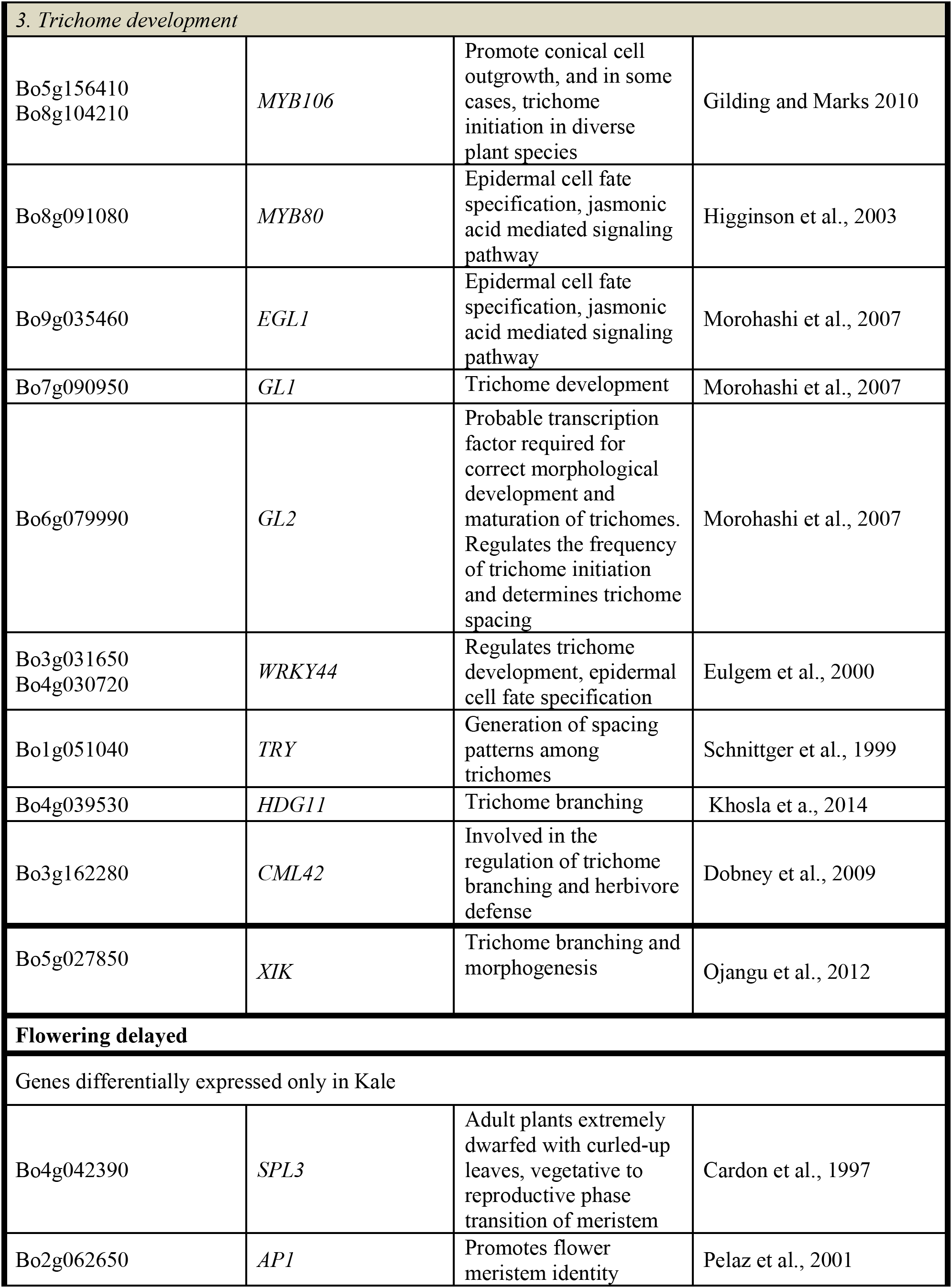
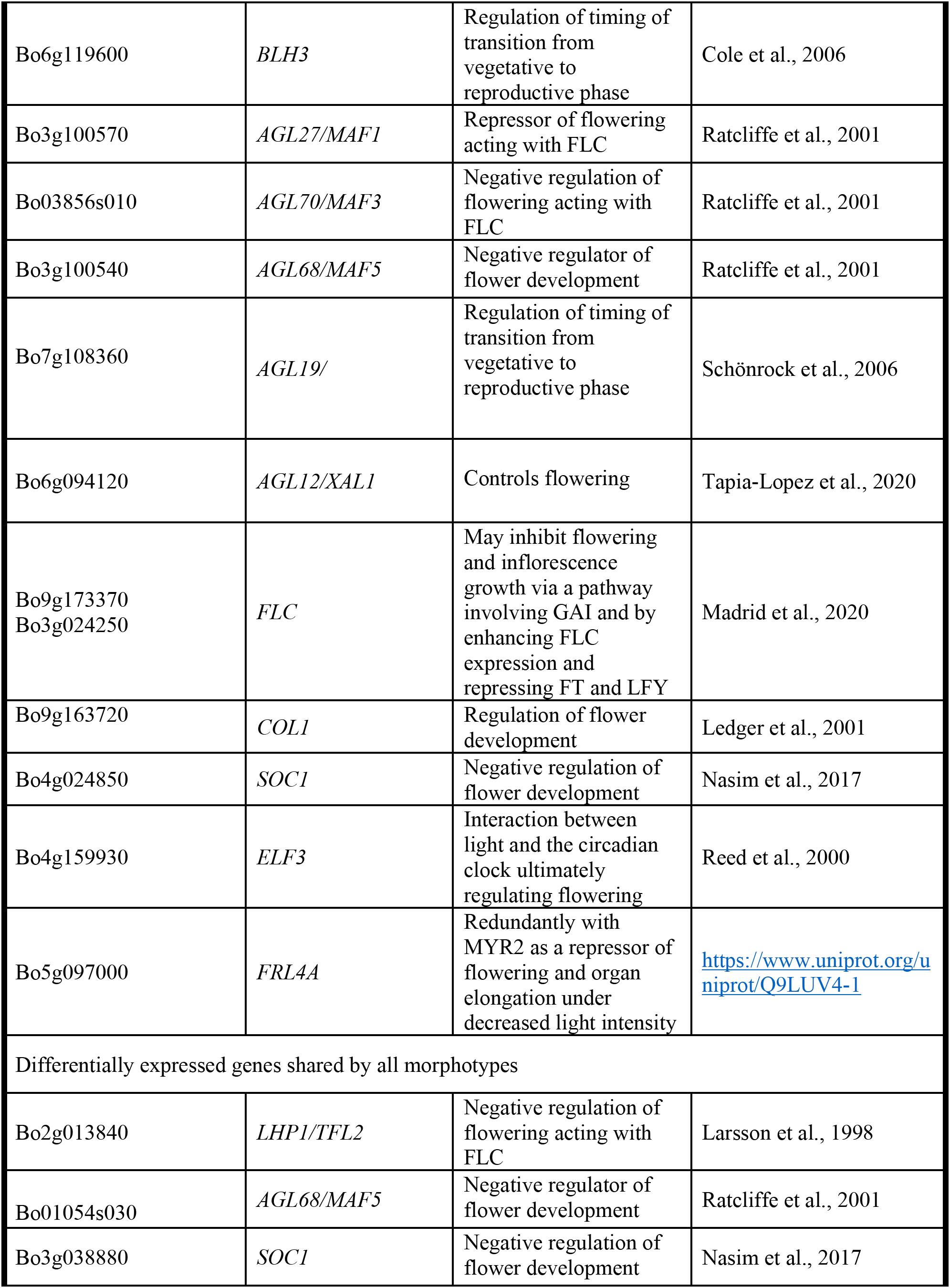
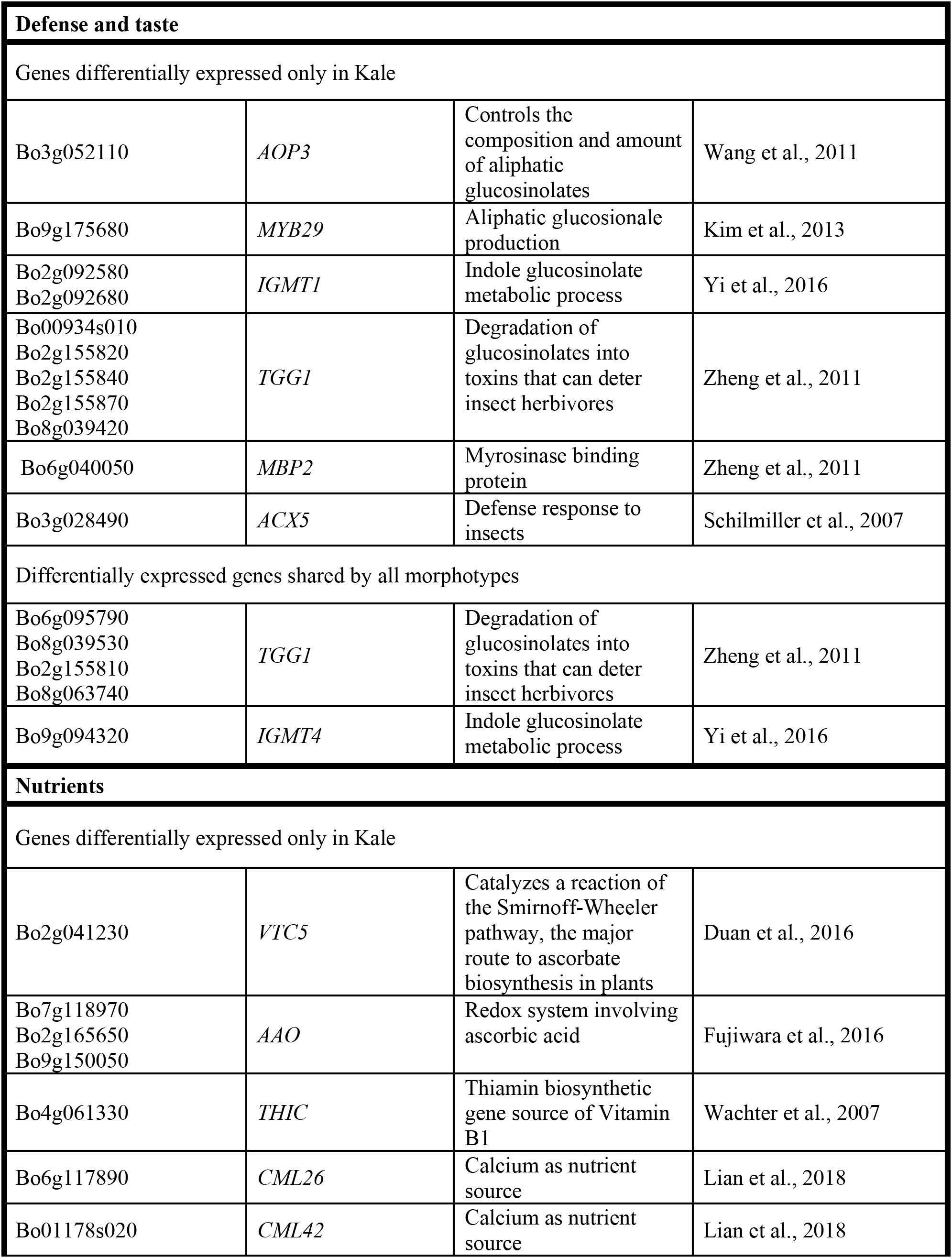
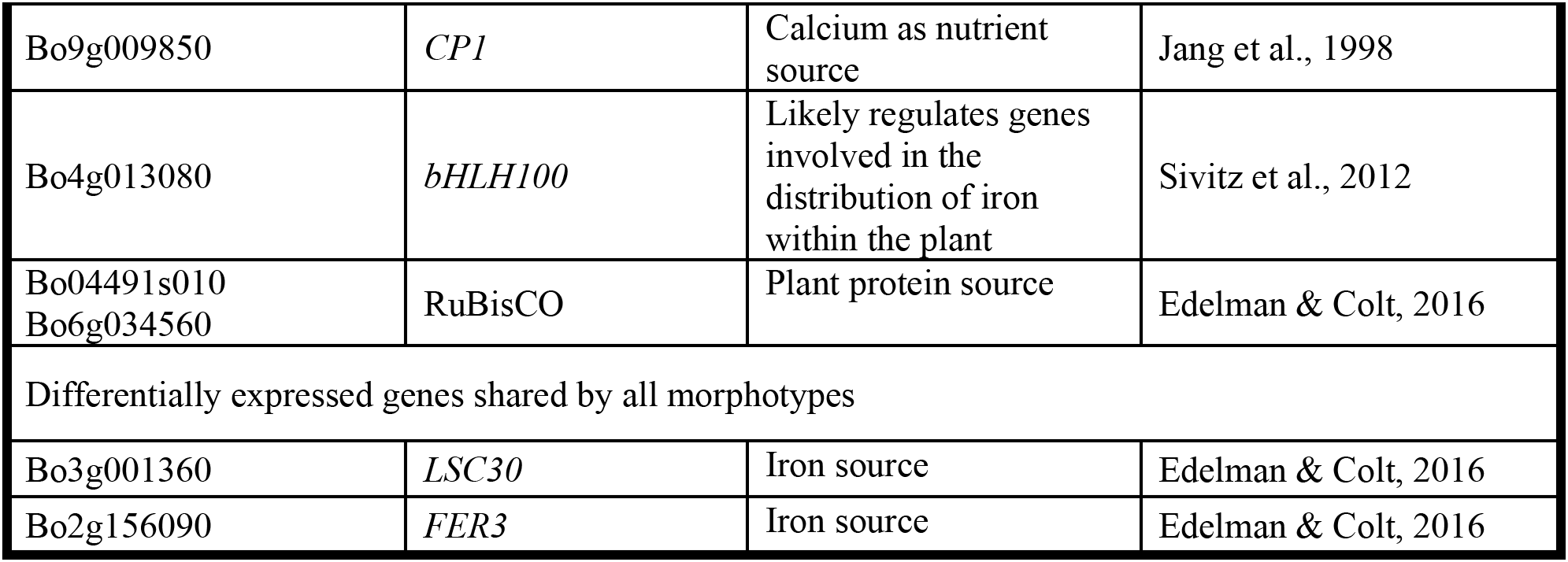
Candidate genes that are differentially expressed in curly kale (red winter kale) and shared among the three morphotypes. Genes involved in plant development, flowering, plant defence responses and taste, nutrition.

#### Leaf development

Differentially expressed genes in kale include two upregulated copies of *AXR1* (Bo8g052710; Bo8g102450). The downregulated LOB domain-containing protein *ASL6* (ASYMMETRIC LEAVES 2-like protein 6) (Bo7g041000) (Semiarti et al., 2001). These genes could have a major role in the highly lobed margin kale phenotype (Fig. 5A). Genes differentially expressed in all three pair-wise comparisons that were potentially involved in the kale domestication syndrome (Fig. 3) include: *PID* (Bo4g182790) that regulates auxin flower and organ formation (Bennett et al., 1995), and *HAT5* (Bo5g155120) which is involved in leaf formation and has been characterized as involved in leaf margin development (Aoyama et al., 1995).

Identified transcription factors included: the AP2-like ethylene-responsive *ANT* (Bo3g153310). This gene is a positive regulator of cell proliferation, expansion and regulation of ectopic meristems (Shigyo et al., 2006), matching what was observed in the kale phenotypes (Fig. 2Q). The growth-regulating factors 3, 4 (*GRF3* and *GRF4*; Bo4g186220 and Bo4g122960) involved in the regulation of cell expansion in leaf and cell proliferation and the Basic helix-loop-helix protein 64 *AtbHLH64* (Bo3g094060), an atypical *bHLH* transcription factor were also identified here (Fig. 5A; Suppl. Table 21).

A significant amount of trichomes in kale leaves were observed during development (Fig. 2 N, H). More than 10 genes related to trichome formation (Table 1, Fig. 6A) were differentially expressed in kale, some of those included: Two upregulated copies of an Myb-related proteins *MYB106* (Bo5g156410 and Bo8g104210). These genes can act as both a positive or negative regulators of cellular outgrowth and promote trichome morphogenesis and expansion (Gilding and Marks, 2010). A Homeobox-leucine zipper protein GLABRA 2-Like *GL2* (Bo6g079990) was upregulated in kale. This gene is required for correct morphological development and maturation of trichomes, regulates the frequency of trichome initiation and determines trichome spacing in *Arabidopsis* (Morohashi et al., 2007). Two expressed copies of the *WRKY* transcription factor 44 (Bo3g031650 and Bo4g030720), and the transcription factor TRIPTYCHON *TRY* (Bo1g051040) involved in trichome branching, were also differentially expressed in kale, among others (Table1, Fig. 6A).

#### Flowering

Kale phenotypes display a significant delaying in flowering in comparison to other *Brassica oleracea* morphotypes. We hypothesized this maintenance in a vegetative stage, was an important character during domestication due mainly to the harvest of fresh leaves for consumption. Our mature kale phenotypes did not display flowering during the course of our experiments (24 days) (Fig. 1). A series of Agamous-like MADS-box proteins were exclusively expressed in kale (Fig. 3B) including two downregulated genes *AGL12* (Bo6g094120) and *AGL70* (Bo03856s010) and an upregulated *AGL27* (Bo3g100570). MADS-box proteins such a *SOC1* (Bo4g024850) was downregulated, and two copies of FLOWERING LOCUS C (Bo9g173370, Bo3g024250) were upregulated (Table 1, Fig. 5A), both involved in the negative regulation of flowering, together with the MYB-related protein *R1 MYB1R1* (Bo6g119010) (Suppl. Table 21). Other genes differentially expressed in all three pair-wise comparisons that were potentially involved in the negative regulation of flowering (Fig. 3) included: *SCO1* (Bo4g024850, Bo3g038880), *SPL3* (Bo4g042390), *AP1* (Bo2g062650), *BLH3* (Bo6g119600), *AGL68/MAF5* (Bo3g100540) among others (Table 1, Fig. 5A)

#### Plant defense

Differentially expressed genes involved in plant defense exclusively expressed in curly kale leaves (Fig. 3b) include: Two copies of the upregulated Myb-related protein *MYB96* (Bo2g082270) involved in the activation of cuticular wax biosynthesis that could potentially deter herbivores. Five upregulated copies of a glucosinolate synthesis and regulation enzyme Myrosinase *TGG1* (Bo00934s010, Bo2g155820, Bo2g155840, Bo2g155870, Bo8g039420), involved in the degradation of glucosinolates to produce toxic degradation products that can deter insect herbivores (Zheng et al. 2011). Last, two copies of the Indole Glucosinolate O-Methyltransferase, one was upregulated *IGMT1* (Bo2g092580), an a second one downregulated *IGMT1* (Bo2g092680) (Table 1, Fig. 5B), and *MYB29* (Bo9g175680) all involved in plant defense against fungus (Suppl. Table 21). Refer to Table 1 and Fig. 5B more other genes potentially involved in plant taste and defense.

#### Nutritional value of leaves

Kale is known to have a great nutritional value including nutrients like Vitamin C, calcium and iron (Edelman & Colt, 2016). Most notable, were three copies of the L-ascorbate oxidase (*AAO*) (Ascorbase) (EC 1.10.3.3)(Bo2g165650, Bo9g150050, Bo7g118970) involved in Vitamin-C synthesis, all three of which were downregulated in kale (Table 1; Fig. 5B) and the enzyme Ferritin-1 *LSC30* (Bo3g001360) proposed as a source of iron. Refer to Table 1 and Fig. 5B more other genes potentially involved in nutritional value of kale leaves.

## DISCUSSION

The vegetative domestication syndrome of kale is characterized by apical dominance, leaf morphology, maintenance of a vegetative stage for high leaf production, taste/defense, and nutritional value. Differentially Expressed Genes (DEGs) were identified in all pair-wise comparisons. In the cabbage comparisons these were enriched amongst syntenic genes, while being depleted in non-syntenic genes (Fig. 4). This aligns with recent findings that genes derived from the ancient *Brassica* polyploidy are more likely involved in domestication in *B. rapa* (Qi et al., 2019). However, in the kale-TO1000 comparison, no such enrichment was found. As the subgenomes of *B. oleracea* have been previously reconstructed, we further explored the relative contribution of each. The LF (least-fractionated) subgenome contains the most genes and so unsurprisingly also had the most DEGs and was over-represented in the cabbage-TO1000 comparions, however, it was underrepresented in the kale-TO1000 comparison. In contrast the MF1 (more-fractionated 1) subgenome was consistently overrepresented. The MF2 (more-fractionated 2) subgenome was overrepresented in the cabbage-TO1000 comparison. Given the limited number of morphotypes in this study, we cannot conclude a general pattern regarding the relative importance of syntenic vs non-syntenic genes and each subgenome, however, the over-representation of the MF1 subgenome in each comparison should be explored further.

We focused our search on DEGs that were amongst comparisons with kale, reasoning that these were more likely to underly the unique characteristics of kale. We tested for enrichment in GO terms and KEGG pathways to gain a global view of the differences. Many more terms and pathways were enriched amongst higher expressed genes, despite several thousand DEGs being identified in both groups. This may suggest more coordination in the regulation of kale higher-expressed genes or simply better annotation of genes in this list. Enriched terms and pathways in all categories were typically involved in metabolism or defense, while we found no clear enrichment of developmental GO terms. Changes in these functions would affect the taste, texture, and nutrition of leaves and thus likely have been affected by human selection. There was a relative absence of developmentally-related GO terms, with the exception of senescence related terms in the kale-TO1000 comparison. This overall lack of enrichment for developmentally-related GO terms could result from either insufficient annotation of developmental genes or due to more subtle expression differences that would require tissue-specific or single-cell approaches. Alternatively, the large developmental changes observed could be caused by differential expression of only a small number of key genes. To address this, we examined sets of genes, whose function based on homology and previous work, suggest an involvement in the developmental traits examined in this study. This analysis found many differentially expressed candidate genes shared amongst comparisons with kale, supporting their potential role in the kale domestication syndrome.

We identified potential candidate genes for the morphological and chemical differences among *Brassica* morphotypes. First, we found differentially expressed genes that set kale apart in terms of its morphology. Apical dominance has been proposed as a main character involved in crop domestication (Doebley et al., 1997). Suppression of axillary branches aiming to concentrate resources in the main stem is a salient phenotypic character in most *Brassica* crops. Apical dominance during domestication in kale can be explained by the indirect theory of apical dominance as auxin-induced stem growth inhibits bud outgrowth by diverting sugars away from buds since growing stems are a strong sink for sugars (Kebrom, 2017). Here genes involved in apical dominance and branching suppression were differentially expressed (Table 1, Fig. 5).

Kale displays morphological diversity in leaves, including leaf shape, size and margins, trichome development, colors among others which is likely the result of genetic variation, selected by plant breeders. Environmental Scanning Electron Microscopy (ESEM) showed that many of the characteristic leaf traits of kale are established early in leaf development. We collected tissues during kale leaf initiation and primary morphogenesis (the upper portion of the stem including immature leaves, and apical and lateral meristems) to capture candidate genes related to the establishment of basic leaf structure such lamina initiation, specification of lamina domains, and the formation of marginal structures such lobes or serrations. Leaf development encompasses three continuous and overlapping phases. During leaf initiation, the leaf primordium emerges from the flanks of the stem apical meristem (SAM) at positions determined by specific phyllotactic patterns (Fig. 2 M-N). In the second phase, the leaf expands laterally and primary morphogenesis events occur from specific meristematic regions at the leaf margin (blastozones) (Fig. 2A-F). In the third phase of secondary morphogenesis, extensive cell expansion and histogenesis occurs (Fig. 2 G-L, Q, R).

Three types differentially expressed genes related to leaf morphogenesis phenotype in kale were found in our samples (Table 1; Fig. 3 and 5), genes: (1) involved in margin development, (2) involved in cell proliferation, expansion and the development of ectopic meristems and (3) involved in trichome development. The Lateral Organ Boundaries LOB domain-containing protein 4 (Asymmetric Leaves 2-like protein 6) *ASL6* (Bo7g041000) was identified as a potential important gene responsible of the curly kale leaf phenotype (Semiarti et al., 2001) (Fig. 1, Fig. 2E, H, K). Loss-of-function mutations in the *AS2* homolog gene in *Arabidopsis* result in asymmetric leaf serration, with generation of leaflet-like structures from petioles and malformed entire veins and the ectopic expression of class 1 *KNOX* genes in the leaves (Matsumura et al., 2009). *ASL6* was differentially expressed and upregulated in kale. Asymmetric placement of auxin response at the distal leaf tip precedes visible asymmetric leaf growth. A second set of candidates involved two copies of the BEL1-like homeodomain protein 1 *BEL1/BHL1* (Bo3g029830, Bo4g142080) that were found to be over-expressed in kale. This gene has been involved in ovule development (Bellaoui et al., 2001), but since we did not collect any reproductive tissue in our sampling, we suggest this might have some other developmental role in kale.

In some species with simple leaves, *KNOXI* over-expression can lead to variable phenotypes, which include knot-like structures on the leaves, curled or lobed leaves and ectopic meristems on leaves (Tsiantis and Hay, 2003). Our differential expression analysis did not show *KNOXI* over-expression, however, the differential expression of *BELL* genes, another class of TALE HD proteins found in plants, was detected. Yeast two-hybrid studies have indicated that *KNOX* proteins interact with BEL1-like (*BELL*) proteins. The mRNA expression patterns of *KNOX* and *BELL* proteins overlap in meristems, suggesting that they may potentially interact in vivo (Smith et al, 2000; Smith et al., 2002). Analogous *BLH–KNOX* interactions have also been reported in other plants, such as potato (*Solanum tuberosum*) and barley (*Hordeum vulgare*), indicating that these interactions are evolutionarily conserved and that the interaction is probably required for their biological functions (Muller et al., 2001; Chen et al., 2003). A matter of debate is whether *KNOX* and *BLH* can exert some of their functions independently of each other or whether the formation of *KNOX/BLH* heterodimers is mandatory for TALEs to work. We did not find differential expression of *KNOX* genes in this analysis perhaps for several reasons: (1) *KNOX* do not participate in lobes formation in *Brassica oleracea*; (2) we did not target the correct point in development to capture *KNOX* expression with relation to leaf lobation. Perhaps these genes are expressed later during development in kale leaves, however we collected tissues from early leaflets where lobes are starting to develop; (3) a change in *BELL* expression is sufficient to affect *KNOX* and *BELL* interactions and alter leaf morphology without a corresponding change in *KNOX* expression, if they were functionally redundant then *KNOX* expression would suffice to hide any effects of *BELL* expression differences. In some species, such as legumes dissected leaves do not express *KNOX* genes; on the contrary *KNOX* expression has been observed in leaf primordia of species with un-lobed leaves, such as *Lepidium oleraceum* (Piazza et al., 2010). We have also found several downregulated Growth-Regulating Factors (*GRF3* and *GRF4*) that have been shown interact with KNOX genes (Kuijt et al., 2014). A third gene involved in cell proliferation, expansion and the development of ectopic meristems included the NGATHA *NGA1* (Bo3g039420) involved in cell proliferation (Lee et al., 2015), however it has been reported as downregulated for cell over proliferation and in our copies was upregulated (Table 1).

Epidermal cells are stimulated to differentiate into trichomes when a regulatory complex including transcription factors are triggered (Yang and Ye, 2013). ESEM results showed hairiness and individual trichomes are seen in kale leaf blades (Fig 2 N, Q). Differentially expressed genes involved in trichome formation included two *MYB* transcription factors (Higginson et al, 2003; Gilding and Marks, 2010; Pattanaik et al., 2014), the enhancer of *GL3 EGL3* (Bo9g035460), the Homeobox-leucine zipper proteins *GL1* (Bo7g090950) and *GL2* (Bo6g079990) (Morohasi et al., 2007), the transcription factor *TRY* (Bo1g051040) (Schnittger et al., 1999) and homeodomain GLABROUS 11, *HDG11* (Bo4g039530) (Khosla et a., 2014). These five genes either interact or act redundantly to promote trichome differentiation and were upregulated in kale (except *HDG11*) (Table 1; Fig. 5A). Trichome regulatory genes have been studied in *Brassica villosa*, a wild C genome relative *B. oleracea* that is densely covered by trichomes. Nayidu et al. (2014) found that *TRY* was upregulated in trichomes of *B. villosa* in contrast to *Arabidospis* and other *Brassica* species where this gene has been proposed as a negative regulator. Here we found *TRY* was upregulated in leaves and meristems of kale in comparison to *ELG3*, *GL1*, *GL2* (Table 1; Fig. 5A).

Another set of candidate genes for the kale domestication syndrome we looked for, are floral transition genes. Floral transition is a major development switch controlled by regulatory pathways, integrating endogenous and environmental cues (Koornneef et al., 2004). We suggest that during kale domestication, delaying of flowering was necessary to maintain leaf production as main consumable. Four main regulatory pathways directly or indirectly involved in flowering have been described in *Arabidopsis thaliana*: photoperiodic, autonomous, vernalization, and gibberellin (Okazaki et al., 2007). The central integrator of flowering signals is the Flowering Locus C (*FLC*) that encodes a transcription factor of the MADS box family. While *A. thaliana* contains only one *FLC* gene, five copies have been described for *B. oleracea* (Okazaki et al. 2007). We found two copies of this gene being differentially expressed in kale. One was upregulated (Bo9g173370) and together with Agamous-like MADS-box protein *AGL27/MAF1* (Bo3g100570) have been proposed as floral repressors (Ratcliffe et al., 2001). A second copy of the *FLC* (Bo3g024250) gene was found to be shared by the three morphotypes kale, cabbage and TO1000 (Table 1, Fig. 5A).

Pathways to domestication in kale may involve selection of plants with the right balance of phytochemicals concentrations. The direction of selection solely towards a reduction in phytochemicals through time was not the case in kale since persistence of pungency and bitterness could have reduced competition from mammals as well as loss to pests, during cultivation and storage (Deham et al., 2020). In *Brassica* species, most glucosinolates are biosynthesized from methionine. A series of copies of the Myrosinase enzyme *TGG1* were differentially expressed and upregulated in kale (Bo00934s010, Bo2g155820, Bo2g155840, Bo2g155870, Bo8g039420) (Zheng et al., 2011). This enzyme breaks down glucosinolates into toxins (isothiocyanates, thiocynates, nitriles, and epithionitriles) upon leaf tissue damage. Therefore, glucosinolates may act as a potent feeding deterrent for generalist insect species, as their toxicity causes developmental and fitness damage (Santolamazza-Carbone 2014, Yi et al., 2015). Gene ontology terms related to glucosinolates and response to herbivores were also found enriched in our analysis specifically in the comparison between kale vs. cabbage terms.

Several genes were identified related to nutritional content in kale, for example precursors and enzymes related to vitamin C, GDP-L-galactose phosphorylase 2 *VTC5* was upregulated in kale (Bo2g041230) (Duan et al., 2016) and several copies of L-ascorbate oxidase (*ASO*) (Ascorbase) (EC 1.10.3.3) *AAO* (Bo2g165650, Bo9g150050, Bo7g118970) (Fujiwara et al., 2016) were downregulated. Precursors of the vitamin B1 Phosphomethylpyrimidine synthase chloroplastic (EC 4.1.99.17) *THIC* (Bo4g061330) (Wachter et al., 2007). The RuBisCO large subunit-binding protein subunit alpha chloroplastic (60 kDa chaperonin subunit alpha (Bo04491s010) and large subunit-binding protein subunit beta chloroplastic (60 kDa chaperonin subunit beta) (Bo6g034560) (Edelman & Colt, 2016). Differentially expressed genes found in the three morphotypes included Ferredoxin-1 chloroplastic (AtFd1) *LSC30* (Bo8g059760) (Table 1, Fig. 5B).

## CONCLUSIONS

This study provided leaf developmental analysis and transcriptomics for *Brassica oleracea* morphotypes kale, cabagge and TO1000, three economically important Cole crops with distinct plant morphologies and domestication syndromes. RNA-Seq experiments allow the discovery of novel candidate genes like those involved in domestication syndromes. We identified gene expression patterns in *Brassica oleracea* that are shared among two vegetative morphotypes cabbage and kale, and the “rapid-cycling” morphotype TO1000, and estimated the contribution of morphotype-specific gene expression patterns that set kale apart. During kale domestication farmers could have been selecting for different flowering times, taste and herbivore defense from plants in the wild that have different vernalization times, different taste, resistance to herbivores or that flower at different times during development and all variations in forms could have arisen from that.

## Acknowledgements

The authors thank G. Conant, J. Birchler, M. Liscum, C. Henriquez, M. Tang, A. Reyes and reviewers for valuable comments on the manuscript. The authors acknowledge the following research grants and granting institutions: National Science Foundation (DEB 1146603, DEB 1209137), Ministerio de Ciencia y Tecnologia de Colombia (64777) and The Newton Fund, UK-Colombia mobility grant.

## Author Contributions

TA: Co-developed questions and framework, obtained funding, made collections, performed microscopy analysis, growth chamber experiments, lab work including library sequencing preparation, performed analyses, wrote manuscript. CH: Performed analyses, performed lab work, wrote and edited manuscript. PM: Co-developed questions and framework, provided funding and mentored students. JCP: Co-developed questions and framework, provided funding and mentored students.

## Data availability

Raw reads and assemblies have been deposited in NCBI: https://www.ncbi.nlm.nih.gov/geo/query/acc.cgi?acc=GSE149483

Additional data can be found in Figshare (https://figshare.com) (Suppl. Tables 2-21): https://doi.org/10.6084/m9.figshare.13224383xs

Commands and their main arguments used for the main steps of analyses: https://github.com/niederhuth/The-molecular-basis-of-Kale-domestication

